# The genomic landscape of asymmetric introgression tracks chromosomal characteristics in tinamous (Aves: Tinamidae)

**DOI:** 10.1101/2025.11.26.690533

**Authors:** Lukas J. Musher, Gregory Thom, Therese A. Catanach, Thomas Valqui, Robb T. Brumfield, Alexandre Aleixo, Kevin P. Johnson, Jason D. Weckstein

## Abstract

We use single genomes from seven tinamou species (Palaeognathae: Tinamidae) and apply a genome-architecture-aware phylogenomic approach to (1) quantify the landscape of introgression, (2) discriminate among conflicting genealogical signals in the genome, and (3) reconstruct evolutionary history. Using summary statistics and full likelihood, we detect pervasive asymmetric introgression between two sympatric Amazonian species clouding phylogenetic relationships in this group. However, these introgression results are sensitive to the assumed species tree. When assuming the Z-chromosome phylogeny as the species tree, the landscape of introgression is non-random; introgression is negatively associated with chromosome length, elevated at macrochromosome ends, and near zero for the Z-chromosome (the avian sex chromosome). A pseudo-autosomal region on the Z-chromosome, which recombines homologously with that of the W-chromosome, shows elevated introgression comparable to that of autosomes. Our results resolve the species tree of tinamous and imply a history of linked selection purging migrant alleles that are deleterious in the hybrid genome. This selection is countered in regions of the genome expected to experience elevated recombination, allowing introgressed alleles to accumulate across >20% of the genome. This work demonstrates a genome-architecture-aware approach to resolving the species tree in the face of introgression. As chromosome-level reference genomes spanning the tree of life become increasingly available, phylogenomic studies would benefit from the adoption of genome-architecture-aware approaches like ours. This is because the genomic landscapes of introgression and genealogy are largely predicted by chromosomal characteristics.

## Introduction

Rapidly diversifying clades offer windows into the processes involved in the formation of species, including hybridization and the generation of postzygotic barriers to gene flow (Malinsky et al. 2015; Han et al. 2017; Nelson et al. 2021; Gyllenhaal et al. 2025). However, when diversification happens over a short period of time, genomic processes of incomplete lineage sorting (ILS) and introgression lead to reticulate genomic histories that complicate biologists’ ability to accurately model the historical sequence of speciation events (the species tree), making inferences about speciation itself challenging (Mallet et al. 2016; Bravo et al. 2019; Smith et al. 2025). Differentiating among conflicting genome-wide signals of ILS, introgression, and the species tree is thus crucial for a deeper understanding of the processes driving biodiversity generation (Edelman et al. 2019; Rivas-González et al. 2023; Musher, Del-Rio, et al. 2024).

The phylogenetic signals of ILS, introgression, and the species tree are expected to vary non-randomly across the genome. Many studies have found that the phylogenetic signal for the species tree is elevated in low-recombination regions of the genome and on sex chromosomes, whereas signals of introgression, which track the history of interspecific hybridization, are enriched in high-recombination regions of the genome (Edelman et al. 2019; Martin et al. 2019; Foley et al. 2024; Stiller et al. 2024). This is because, when migrant alleles are incompatible with the local environment or recipient genome, selection may purge neutral or adaptive variation that is linked to maladaptive sites (Schumer et al. 2018). However, recombination counters this effect by breaking up the linkage between maladaptive alleles and neighboring loci such that regions of the genome with elevated recombination should also retain a higher portion of introgressed alleles (Lohmueller et al. 2011; Veller et al. 2023). Although accurately estimating recombination rates across the genome with phylogenomic-scale datasets and small sample sizes is challenging, this issue can be mitigated by using recombination rate correlates to distinguish among discordant phylogenetic signals in the genome. For example, (1) macrochromosomes (>25Mb) have lower recombination than microchromosomes (<25Mb) (Tigano et al. 2022), (2) the distal ends of macrochromosomes have more recombination than centers, and this is independent of centromere position (Haenel et al. 2018), and (3) the major sex (Z or X) chromosome has lower recombination than autosomes (Charlesworth et al. 1987; Irwin 2018). In addition, recombination covaries with other genomic characteristics, such as GC content, because long-term recombination rate variation can shape base composition via GC-biased gene conversion (Singhal et al. 2015). Thus, to discriminate among conflicting genealogical signals in the genome, chromosomal characteristics may be used as a proxy for recombination when limited genomic data precludes direct estimation of recombination rates (Musher, Del-Rio, et al. 2024).

Here we implement a genomic-architecture-aware approach to model the evolutionary history of a clade of seven tinamou (Palaeognathae: Tinamidae) species in the genus *Crypturellus* to disentangle historical genomic signals of ILS, introgression, and speciation. This clade evolved in under 5 million years and is marked by rapid differentiation and potential introgression between non-sister taxa. Our previous work utilized orthologous sequences spread across the whole genome to reconstruct the species tree in this group but could not resolve the relationships within *Crypturellus*, even with tens of thousands of markers (Musher et al. 2025). However, we identified three potential species trees (Figure 1A: T1, T2, & T3) that differed only in the placement of a single species, *C. atrocapillus,* and argued that pervasive genome-wide introgression produced substantial phylogenetic conflict that hampered species tree inference. Here, to further investigate this system, we ask three questions: first, how are signals of genealogy and introgression distributed across the genome? Second, can we utilize information about genome architecture and the landscape of introgression to make robust inferences about the species tree in this clade? And finally, what does this approach reveal about the history of and genomic resistance to hybridization during avian speciation?

**Figure 1:**
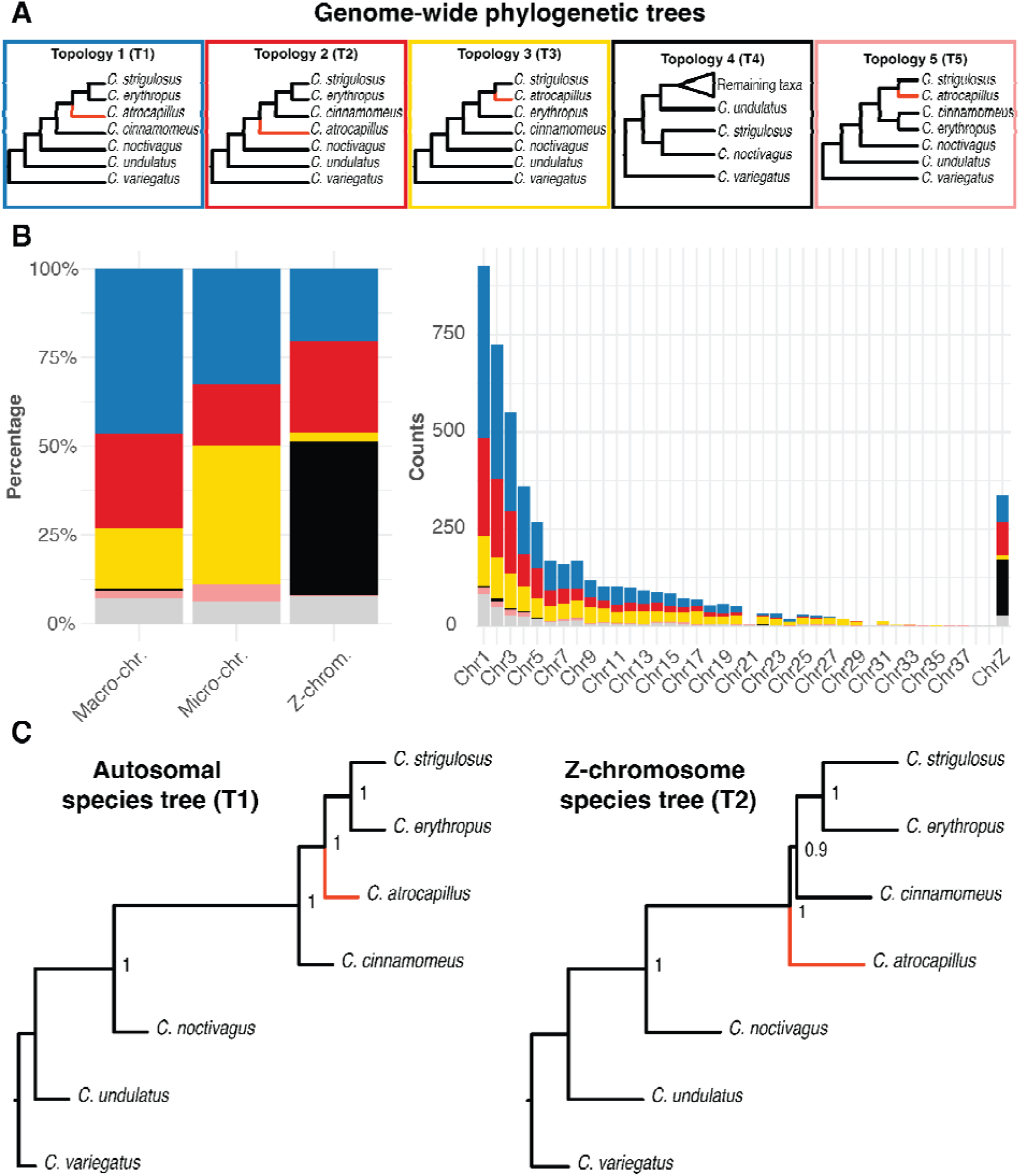
Evolutionary history showed great variation across the genome. **(A)** The top row of boxes show the four most frequently recovered phylogenetic topologies across the genome that differed by the location of *C. atrocapillus* (red branches). (B) These gene trees varied in frequency both within and among chromosomes. T1–T3 were recovered in a previous study as equally supported based on both ultraconserved elements (T1 & T2) and protein coding sequences (T3). (C) Species trees inferred from autosomal and Z-chromosome gene trees differed, recovering T1 for autosomes and T2 for the Z-chromosome.

## Results

### Gene tree heterogeneity is high across the genome

We reconstructed phylogenetic trees for nonoverlapping windows across the genome and recovered five primary topologies when analyzing large (200-kb) windows. These included the previously recovered T1, T2, and T3 (Musher et al. 2025), as well as a new topology primarily recovered from Z-chromosome windows (T4) and another recovered primarily from autosomal windows (T5) (Figure 1B). The most common topology found across autosomal windows was T1 (42.9% of gene trees) but for the Z-chromosome windows, it was unexpectedly T4 (43.2%), which dubiously recovered a sister relationship between the only two female samples in the dataset, *C. strigulosus* and *C. noctivagus* (Figures 1B and 2). The second most common Z-chromosome topology was T2 (26%). Signal for these topologies was not randomly distributed across the genome, with an overrepresentation of T3 from microchromosomes, the distal tips of macrochromosomes (chromosomes 1–8), and at one end of the Z-chromosome.

**Figure 2:**
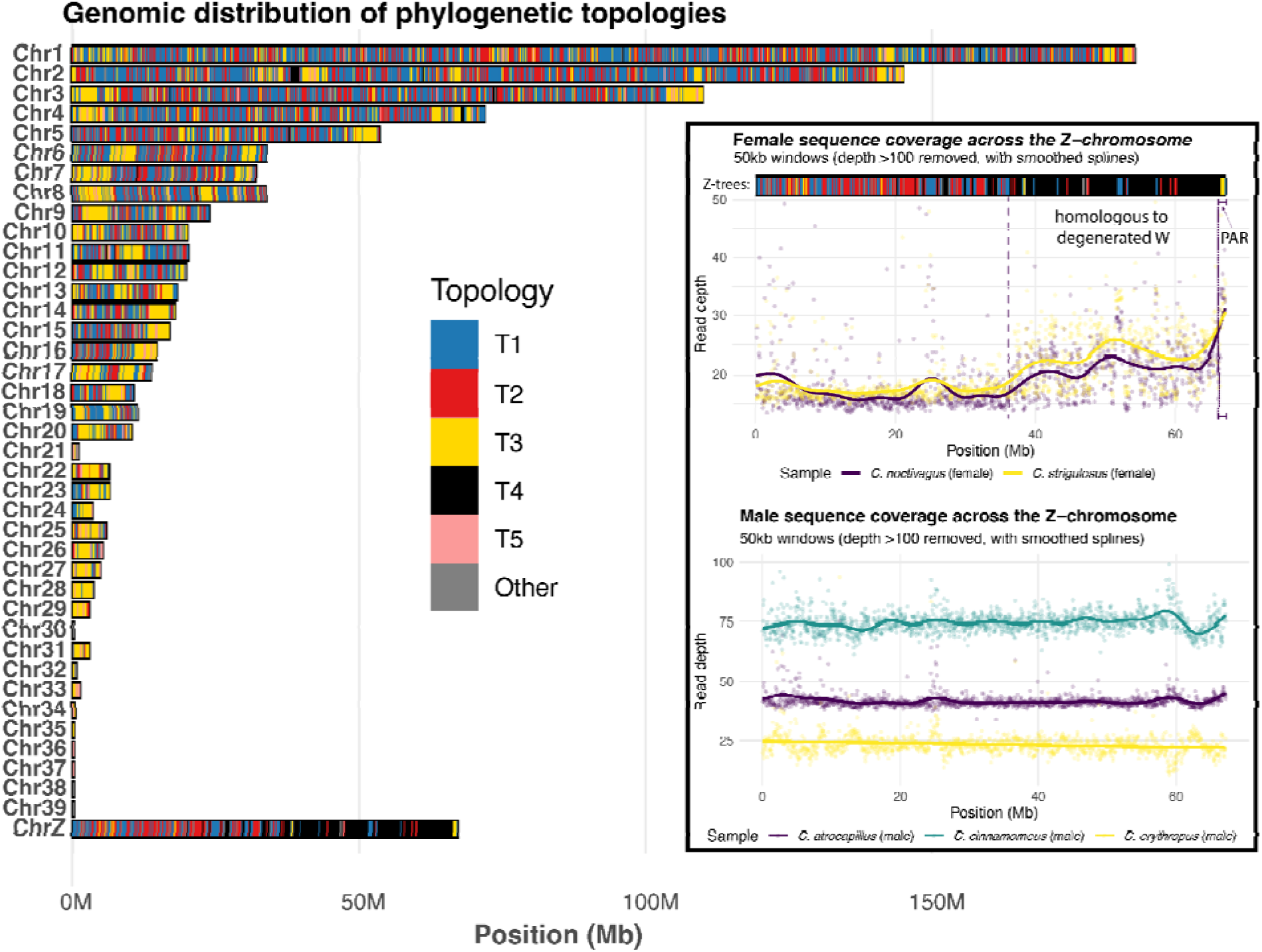
The distribution of tree topologies across all chromosomes in the genome. The bar plot at the left shows the positions of recovered gene tree topologies across each chromosome. Each vertical bar on each chromosome represents a 200 kb window. Colors match the topologies introduced in Figure 1, including T1 (blue), T2 (red), T3 (yellow), T4 (black), and T5 (pink). Other topologies are not parsed further and are shown in gray. The inset box shows how changes in read depths for female samples illuminate the pseudo-autosomal region (PAR) and reveal mapping error on the Z-chromosome. The top scatterplot shows read depth across the Z-chromosome for two female (heterogametic) samples in the dataset, *C. strigulosus* and *C. noctivagus*. At the top of the plot, we also show gene tree topologies across the Z-chromosome from Figure 2, indicating that the region of the Z-chromosome overrepresented by T4 gene trees corresponds to a region of the Z-chromosome that maintains high sequence similarity with the W-chromosome. The PAR is located at the end of the Z-chromosome, which is marked by roughly twice the depth of the first half of the Z. The bottom scatter plot shows uniform depths across the Z-chromosome consistent with expectations of male (homogametic) samples.

### Autosomal and Z-chromosome species trees differ

To assess broad-scale evidence for the species tree from genome-wide data, we analyzed autosomal and Z-linked data separately using a multi-species coalescent (MSC) approach in ASTRAL-III (Zhang et al. 2018). MSC methods for estimating the species tree assume input gene trees are from non-recombining genomic loci. We therefore used smaller windows of 10 kb for estimating the species tree. The inferred autosomal MSC tree matched the phylogenetic topology T1, recovering *C. atrocapillus* as sister to *C. strigulosus* + *C. erythropus*. The inferred Z-chromosome MSC tree matched topology T2, recovering *C. atrocapillus* as sister to the clade containing *C. strigulosus*, *C. erythropus*, and *C. cinnamomeus* (Figure 1C). These results are in line with other studies, which commonly recover unique autosomal and sex-chromosome evolutionary histories. Specifically, many studies have shown that the signal for the species tree is often overrepresented on sex-chromosomes (Edelman et al. 2019; Martin et al. 2019; Foley et al. 2024; Stiller et al. 2024).

### The landscapes of genealogy and introgression track chromosomal characteristics

We measured levels of introgression across the genome using a summary statistic *f_dM_* (Malinsky et al. 2015), an extension of the D-statistic (Patterson et al. 2012) that quantifies the proportion of introgression assuming a species tree. As genome architecture is often predictive of the landscape of introgression, we hypothesized that levels of introgression should covary with chromosomal characteristics only when assuming the correct species tree. We therefore modeled *f_dM_* twice: once assuming the autosomal tree (T1; positive *f_dM_* indicates introgression between *C. cinammomeus* and the branch containing *C. strigulosus* + *C. erythropus*) and once using the Z-chromosome tree (T2; positive *f_dM_*indicates introgression between *C. atrocapillus* and the branch containing *C. strigulosus* + *C. erythropus*). Consistent with our hypothesis, we found that levels of introgression varied across the genome and tracked chromosomal characteristics when assuming the Z-chromosome tree (T2), but not the autosomal tree (T1; Figure 3). Specifically, we found weak or absent covariation between genome architecture and *f_dM_* when assuming T1 as the species tree (Figure 3A). However, when assuming T2 (the Z-chromosome tree), patterns of introgression tracked *a priori* expectations associated with genome architecture. Specifically, (1) macrochromosomes had lower introgression than microchromosomes, (2) there was a strong negative association between introgression and chromosome length, and (3) the Z-chromosome showed lower introgression on average than autosomes, nearing zero for most of its length. Dunn tests confirmed that these differences were significant (P<0.0001 for all pairwise comparisons). Finally, we found that macrochromosomes, and some larger microchromosomes, had a significant pattern of increasing introgression toward their tips. This was true when looking at all nine macrochromosomes combined, as well as when looking at the largest chromosomes in the genome individually (Figures 3B & S1). However, when assuming T1 as the species tree, these patterns were typically counter to *a priori* expectations or showed weaker relationships (Figures 3A, S2).

**Figure 3:**
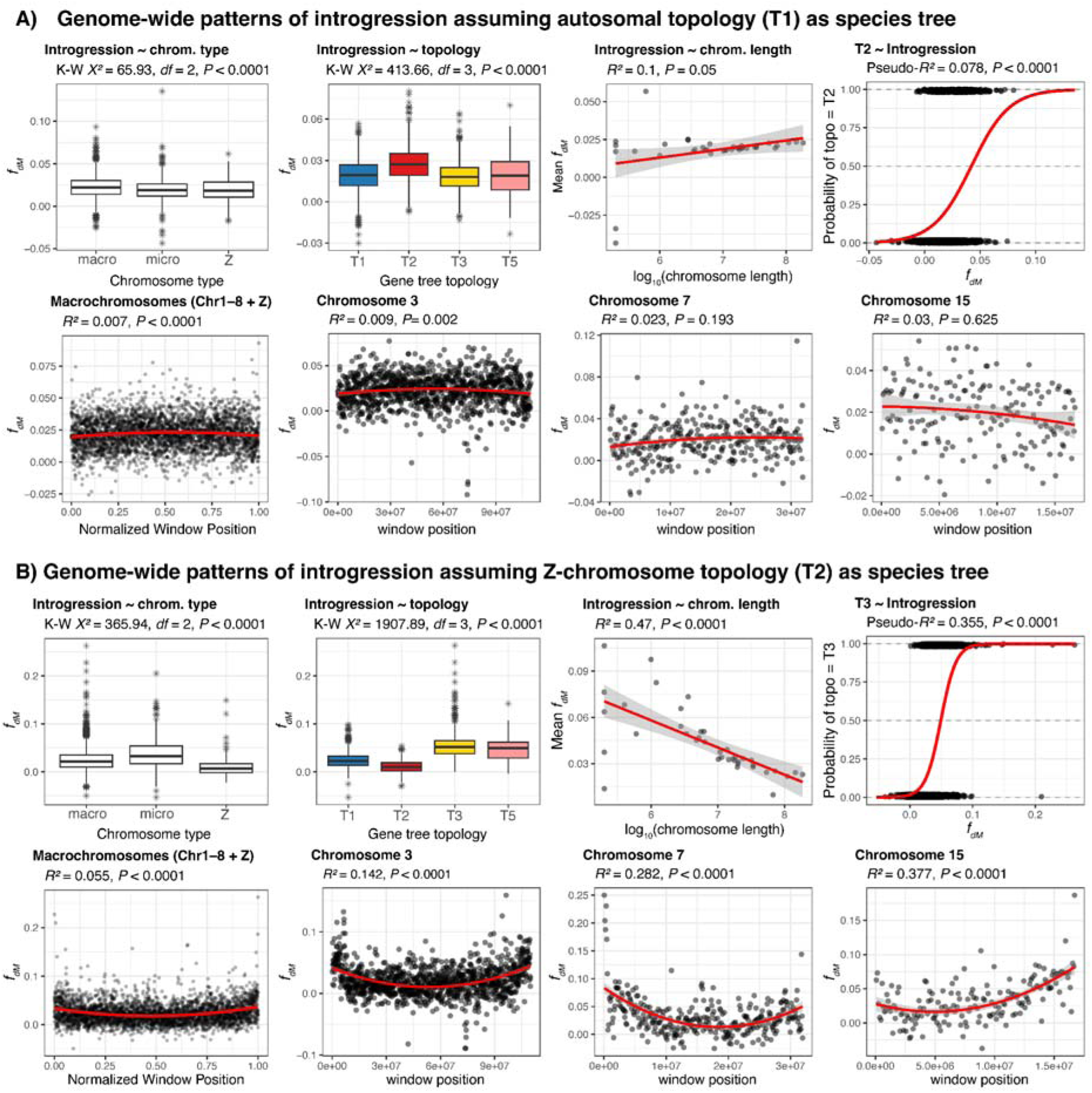
Levels of introgression across the genome covary expectedly with chromosomal characteristics when assuming T2, but not T1, as the species tree (erroneous Z-chromosome windows removed). (A) Patterns of genome-wide introgression assuming T1 (the autosomal topology) as the species tree. Top row (from left to right): differences in levels introgression between macro-, micro-, and Z-chromosomes, differences in introgression for windows recovering alternate topologies T1–T3 and T5, covariation between chromosome length and introgression, and logistic regression showing the probability of a window recovering T2 as a function of introgression. Bottom row: quadratic models relating introgression by normalized position on macrochromosomes, and actual position for three example chromosomes (Chr. 3, 7, and 15). Positive values of *f_dM_* indicate introgression between *C. cinnamomeus* and the branch containing *C. strigulosus + C. erythropus.* (B) Patterns of genome-wide introgression assuming T2 (Z-chromosome topology) as the species tree. Plots are the same as in (A) except that the logistic regression model (top right) shows the probability of recovering T3 as a function of introgression. Positive values of *f_dM_* indicate introgression between *C. atrocapillus* and the branch containing *C. strigulosus* + *C. erythropus*.

When assuming T2 as the species tree, the landscape of introgression also covaried with genealogical history. A Kruskal-Wallis test showed that windows recovering different topologies also varied in levels of introgression (*X^2^* = 2068.6, *df* = 4, *P* < 0.0001; Figure 3B). A Dunn Test indicated that windows recovering T3 and T5 each had higher introgression than windows recovering other topologies (*P* < 0.0001), and those recovering T1 had higher introgression than those recovering T2 (*P* < 0.0001). We also used logistic regression to test for a relationship between *f_dM_* and the probability of recovering individual topologies and detected a strong positive relationship between *f_dM_* and the T3 topology (Ψ = 5 × 10^35^, Psuedo-*R^2^* = 0.355, *P* < 0.0001; Figures 3, S3, & S4). T1 was also associated with introgression, but this relationship was weaker and negative (Ψ = 1.1 × 10^5^, Psuedo-*R^2^* = 0.012, *P* < 0.0001). On the other hand, T2 was strongly associated with lower levels of introgression, showing a negative association with *f_dM_* (Ψ = 4.8 × 10^-30^, Pseudo-*R^2^* = 0.196, *P* < 0.0001).

We also found that patterns of genome-wide GC content, expected to covary with recombination rates, mirrored those of introgression (Figures S5 & S6), and that introgression was positively correlated with both gene (*R^2^* = 0.025, *P* < 0.0001) and GC (*R^2^*= 0.051, *P <* 0.0001) content in the genome. Although genes are expected to be targets of selection that might resist introgression, in birds, crossover sites are tied to gene promoters. Thus, regions with high gene content may also experience increased local recombination rate, reducing impacts of linked selection in gene- and GC-rich regions (Singhal et al. 2015). Our results are therefore consistent with a scenario where selection against hybrids purges foreign alleles from the hybrid background most efficiently in low-recombination portions of the genome.

### Introgression was asymmetric

To confirm introgression inferences derived from summary statistics, we applied a full likelihood approach using a multi-species coalescent with migration (MSC-M) (Flouri et al. 2023) model on a subset of 10,000 randomly sampled 10kb windows across the genome. This model confirmed pervasive introgression between *C. atrocapillus* and *C. strigulosus* with much higher migration from *C. strigulosus* to *C. atrocapillus* (ω median = 6.08; 95% Credible Interval [CI] = 5.71–6.46) than from *C. atrocapillus* to *C. strigulsosus* (ω median = 0.024; 95% CI = 0.0025–0.089), indicating introgression was asymmetric. This asymmetry may be driven by significantly higher effective population size (N_e_) of *C. strigulosus* (N_e_ median = 8,128,637; 95% CI = 7,046,446–9,397,264) than *C. atrocapillus* (N_e_ median = 456,575; 95% CI = 78,668–1,210,671). Finally, the MSC-M model estimated three consecutive and rapid divergences in this clade. The first divergence between *C. atrocapillus* and its sister clade occurred ~3.7 mya (95% CI = 3,711,050–3,749,700 mya), with a subsequent split from *C. cinammomeus* at ~3.13 mya (95% CI = 3,117,050–3,139,850) and the most recent split between *C. erythropus* and *C. strigulosus* at ~3.12 mya (95% CI = 3,106,800–3,130,900).

### T4 is associated with putative mis-alignment of W-chromosome reads to a homologous portion of the Z-chromosome

We aimed to discriminate phylogenetic signals of ILS, introgression, and the species tree from those potentially reflecting error by characterizing genome architecture and related variation in recombination rate. The avian sex chromosome, chromosome Z, is expected to have reduced recombination compared with autosomes. However, chromosome Z also harbors a pseudo-autosomal region (PAR), which recombines homologously with that of chromosome W. In paleaognathous birds, the PAR is typically quite long compared to other avian groups, but is reduced in some tinamou species (Xu et al. 2019; Wang et al. 2022). To identify the PAR, which is diploid in the heterogametic sex, we quantified per-base sequence coverage across the Z-chromosome with the expectation that females should have ~2X relative read depth along the PAR compared with the remaining Z-chromosome, which is haploid in females. For the two female samples, we recovered a pattern of read overrepresentation in the second half of the Z-chromosome corresponding to the location of a long region enriched for the T4 topology (Figure 2 inset). This result was absent in male samples, which exhibited uniform read depth across the Z-chromosome. The increase in read depth for this region in female samples was less than two-fold, indicating the section enriched for T4 is not included in the PAR. Instead, we suggest this region of the Z-chromosome is homologous to a non-recombining region of the W-chromosome, which is proximal to the PAR. This is consistent with other studies indicating that degeneration of the W-chromosome occurred relatively recently in ancestors of *Crypturellus* tinamous, meaning that the Z and W chromosomes maintain high sequence similarity in this region (Wang et al. 2022). We therefore interpret these elevated depths for females to represent the summation of erroneously mapped W-chromosome reads with properly mapped Z-chromosome reads. Only the most distal ~1 Mb of the Z-chromosome had relative read depths indicative of the PAR. The PAR was enriched for the T3 topology, harbored a high proportion of introgressed alleles, and contained high GC content (Figures 2, 4A, S5, & S6).

**Figure 4:**
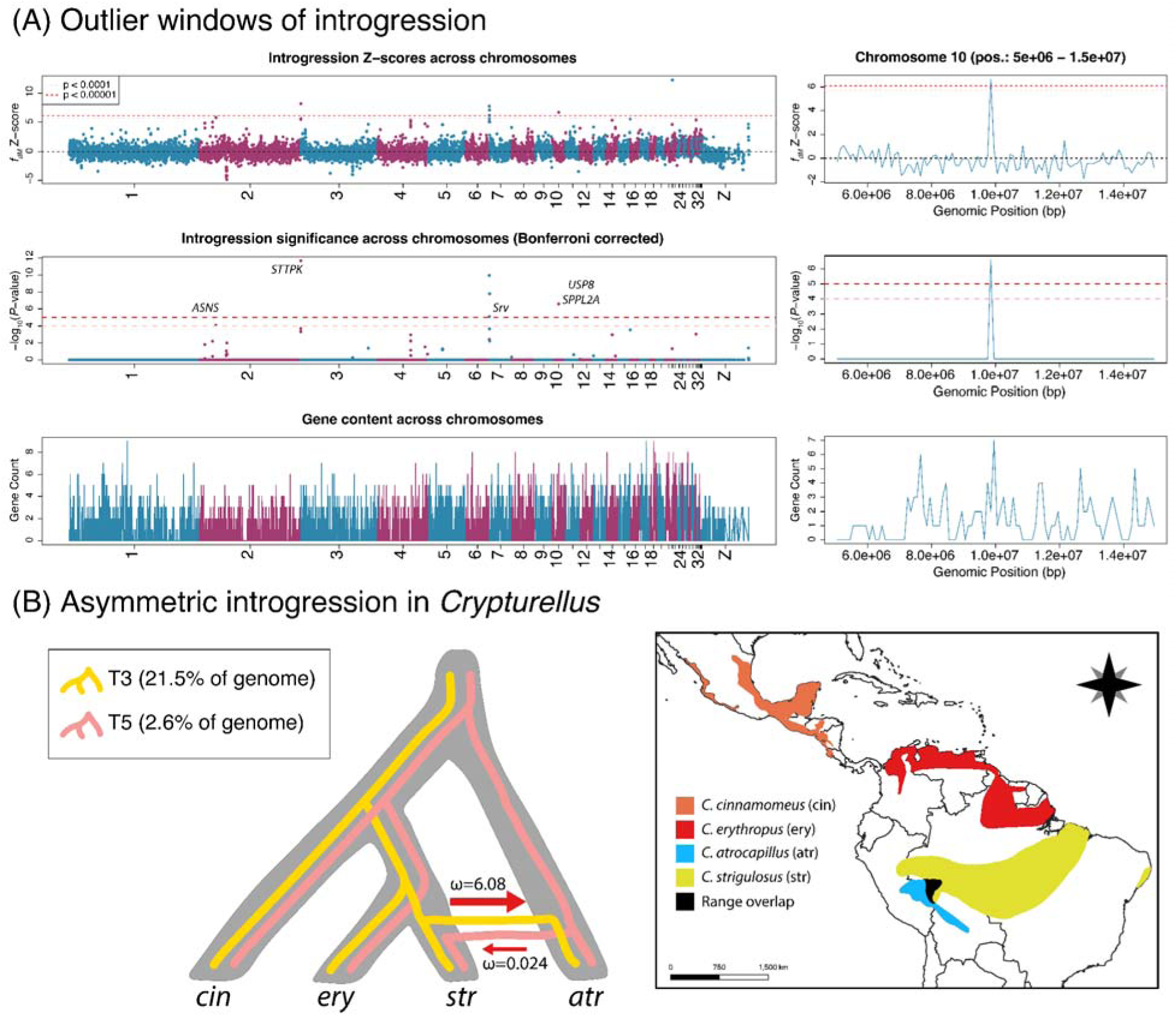
Most windows of significant introgression are located near chromosome ends and lack high gene content except on chromosome 10. (A) Candidate windows of adaptive introgression showing high Z-scores (top), Bonferroni corrected p-values (middle), and gene content (number of genes per 100kb window) for the whole genome (left) and a region of chromosome 10 with high gene content (right). (B) The species tree (T2) is shown in gray with hypothetical T3 (yellow) and T5 (pink) gene trees overlain. The shape and prevalence of the T3 topology compared with the T5 implies that asymmetric introgression from *C. strigulosus* (str) into *C. atrocapillus* (atr) is driving the pervasive signal of positive *f_dM_* across the genome. Mean mutation-scaled migration rates (ω) from the MSC-M model are shown on arrows indicating asymmetric introgression. The map at the right shows the geographic distributions of four species potentially associated with introgression (depending on assumed species tree). Only *C. atrocapillus* (blue polygon) and *C. strigulosus* (yellow polygon) overlap in southwestern Amazonia (black polygon). Range map polygons downloaded from <u>IUCN.org</u>. Tinamou illustrations by TAC.

### Most outlier peaks of introgression occur near chromosomal tips

As a final examination of patterns of introgression across the genome, we identified windows in the genome with measured *f_dM_* that differed significantly from the rest of the genome. Windows with high *f_dM_*contain a high proportion of introgressed alleles that may represent candidates for adaptive introgression, recent introgression, or adaptive cohesion among linked co-evolved genes. We estimated Z-scores for all windows and used a two-tailed permutation test to assign significance*-*values to each window. We identified seven windows with corrected p-values < 0.0001. These occurred on chromosome 2, chromosome 7, chromosome 10, and chromosome 21. Two peaks on different segments of chromosome 2 each contained a gene involved in ATP and nucleotide binding (Serine-threonine/tyrosine-protein kinase [STTPK] and Aspergine synthetase [ASNS]), and a peak on chromosome 7 contained a gene involved in visual perception (7TM GPCR, serpentine receptor class v [Srv]). Overall, each window with significant introgression had relatively low gene content and tended to occur toward the chromosome tips, except on chromosome 10, which had high local gene content and occurred toward the center of the chromosome (Figure 4). The peak on chromosome 10 contained two genes involved in posttranslational modification and protein turnover (ubiquitin carboxyl-terminal hydrolase 8 isoform X1 [USP8] and signal peptide peptidase-like 2A [SPPL2A]). In all, there were 22 genes detected within 500 kb of this high introgression window on chromosome 10 (Table S2).

## Discussion

Here, we demonstrate how using genome architecture as a proxy for the expected landscape of recombination can aid in species tree inference in cases where introgression is pervasive. Specifically, we show that for two sympatric non-sister Amazonian tinamou species, the landscape of introgression covaries with chromosomal characteristics, a proxy for recombination rates. It is now widely appreciated that introgression between non-sister taxa is prevalent in the tree of life (Mallet et al. 2016; Meier et al. 2017; Andersen et al. 2021; Nelson et al. 2021; Musher et al. 2022; Thom et al. 2024). However, inferringing introgression can be challenging in cases where high gene tree heterogeneity clouds species tree inference (Deraad et al. 2025). Indeed, if species’ relationships are uncertain, then there is potential to confound other sources of gene tree discordance (e.g., ILS or technical error) for introgression. Our genome-architecture-aware approach offers a solution to this problem, allowing one to infer the species tree after comparing the inferred landscape of introgression to theoretical expectations based on expected variation in recombination rates across the genome.

A key finding was that the inferred landscape of introgression depended on the assumed species tree when estimating summary statistics, indicating that post-hoc tests of gene flow can be positively misleading if assuming an incorrect species tree. When assuming the Z-chromosome topology as the species tree, we recovered (1) a negative relationship between chromosome length and introgression, (2) increased introgression on microchromosomes and macrochromosome ends, and (3) decreased introgression on the Z-chromosome. These results, when combined with the expectation that signal for the species tree should be enriched on the Z-chromosome, imply that T2 likely represents the species tree for this clade. Because these summary statistics require the user to know the true species tree to make accurate inferences, assessing introgression results in the context of genome architecture may provide a useful check on their application.

Evolutionary theory posits that covariation between introgression and chromosomal characteristics reflects interactions between selection and recombination (Schumer et al. 2018; Martin et al. 2019; Tigano et al. 2022; Thom et al. 2024). Recombination and introgression rates should covary when there is baseline selection against gene flow due to either (1) divergent selection that purges migrant alleles that are deleterious in the local environment (Feder et al. 2012), or (2) selection that purges foreign alleles that cause negative epistatic interactions in a hybrid background (Racimo et al. 2015; Schumer et al. 2018). When recombination rates are low (e.g. on the sex chromosome and centers of macrochromosomes), selection will purge deleterious migrant alleles along with long tracks of neighboring DNA due to genetic hitchhiking (i.e., linked selection), but increased recombination (e.g., on microchromosomes and macrochromosome tips) breaks up the linkage between deleterious alleles and neighboring loci, allowing easier gene flow in these regions of the genome (Li et al. 2019; Martin et al. 2019; Nelson et al. 2021; Tigano et al. 2022).

We also identified seven outlier peaks of introgression (Figure 4A, Table S2). Most of these windows occurred near the tips of chromosomes and could be explained by elevated recombination in these regions coupled with recent introgression events. One peak, however, occurred toward the center of chromosome 10 (Figure 4A). This region harbored 22 genes within 500 kb of the *f_dM_* peak. High gene content might be expected to result in increased positive selection if genes in this region are co-evolved and result in reduced fitness when recombined with alternate alleles. In birds, however, crossover sites are tied to gene promoter regions (Singhal et al. 2015), which may lead to an increased local recombination rate in gene-dense areas, thereby reducing impacts of linked selection. We recovered a positive correlation between gene content and *f_dM_*, which supports this idea. Moreover, in our previous study, species trees reconstructed from coding sequences converged on the T3 (introgression) topology (Musher et al. 2025). Nevertheless, we cannot distinguish scenarios wherein outlier values of *f_dM_*are driven by positive selection on migrant alleles, cohesion among coevolved genes, or recent introgression, each of which could result in a high proportion of introgressed variants. To discriminate among these hypotheses, additional intraspecific sampling is needed to both estimate recombination rates and test for adaptive evolution (Lewanski et al. 2024).

Our results agree with the expectation that introgression should be elevated in regions of the genome that experience elevated crossover rates. Because all chromosomes undergo at least one crossover during Anaphase I of Meiosis (Saito and Colaiácovo 2017), microchromosomes experience more recombination per base than macrochromosomes (Tigano et al. 2022). Moreover, crossing over is initiated at the ends of chromosomes, commonly causing a distal recombination rate bias on large chromosomes but more uniform recombination rates on microchromosomes (Haenel et al. 2018). Although decreased recombination at chromosome centers occurs independently of centromere position (Haenel et al. 2018), recombination rates are further suppressed at centromeres. On most large chromosomes in our dataset, T3 (an introgression topology) was enriched at both tips. However, on chromosomes 5, 6, 15, and 16, T3 was only enriched at one end of the chromosome. We suggest that these four chromosomes may be telocentric or acrocentric (centromeres at or near one end, respectively), however, we could not corroborate this hypothesis based on our short-read data and pseudo-chromosome reference. Although we did not measure selection or recombination, our results are consistent with expectations of these processes driving patterns of introgression. Still, with a relatively distant chromosome-level reference genome and limited intraspecific sampling, we were able to utilize knowledge about genome-architecture to inform phylogenetic inference (Musher, Del-Rio, et al. 2024). We suggest that as chromosome-level reference genomes become increasingly available across diverse taxa in the tree of life, genome-architecture-aware approaches such as should become standard instruments in the phylogeneticist’s toolkit.

### A new case of reticulate Amazonian diversification

Introgression results from biogeographic and genomic processes that affect interspecific interaction and reproductive isolation (Hewitt 1988; Musher 2025). In the Amazon, many phylogeographic and phylogenetic studies are finding evidence of introgression between non-sister species reflecting both biogeographic and genomic processes. For example, many bird and mammal lineages in Amazonia are allopatrically isolated by riverine barriers but experience substantial introgression at narrow headwaters or after rivers move (Moncrieff et al. 2023; Musher, Del-Rio, et al. 2024). In cases where post-zygotic reproductive isolation is weak or absent, the genomes of hybridizing lineages can experience substantial homogenization (Barrera-Guzmán et al. 2022). Here we find that despite more than three million years of divergence, *C. atrocapillus* has accumulated introgressed alleles across >20% of its genome with evidence of polygenic selection against such introgression.

There are four documented examples of hybridization in tinamous, however a recent analysis found that only one of these was convincing (Ottenburghs 2021). Here, we documented a new example of hybridization that has led to genome-wide introgression. Specifically, we detected a consistent signal of introgression between *C. atrocapillus* and the clade that contains *C. strigulosus* and *C. erythropus* marked by positive *f_dM_* across much of the autosomal genome. Both T3 and T5 unite *C. atrocapillus* and *C. strigulosus* as sisters and are associated with significantly higher *f_dM_* than other topologies (Figure 3B). The T3 topology nests *C. atrocapillus* in the clade containing *C. erythropus* and *C. strigulosus*, suggesting that windows recovering this topology contain alleles in the *C. atrocapillus* background, inherited from *C. strigulosus* (Figure 4B). The T5 topology implies the opposite history. Since T3 was nearly ten times more frequent across the genome, the prevalence of this topology indicates asymmetric introgression from *C. strigulosus* into the *C. atrocapillus* genome. Parameter estimation using full likelihood confirmed this finding. This scenario is plausible; unlike other species pairs in this clade, the two species’ distributions overlap in southwestern Amazonia (Figure 4B). Though *C. atrocapillus* is more locally endemic, *C. strigulosus* has a more widespread distribution. Thus, hybridization between these two taxa could result in the asymmetric diffusion of alleles from *C. strigulosus*, the species with a larger effective population size, into *C. atrocapillus*, the species with a smaller population (Lenormand 2002).

Since T3 and T5 are most closely associated with introgression, and T2 the presumed species tree, this raises the question of why T1 is the most common topology across the genome. Although gene trees supporting T1 were associated with higher *f_dM_* values than T2 (Figure 3B), and may be consistent with hybridization between *C. atrocapillus* and the ancestor of *C. strigulosus* + *C. erythropus*, we did not find a strong positive association between this topology and *f_dM_,* instead finding a weak negative association (Figure S4). We suggest that the prevalence of this topology could result from three non-mutually exclusive factors. First, because T1 is associated with higher introgression than T2, it may represent a weak signal of introgression, that leads to clustering of *C. atrocapillus* with the *C. strigulosus/erythropus* clade, but is not strong enough to nest it within the clade (i.e. fewer shared derived alleles). Given that these results are based on relatively long sliding windows, this seems possible since some windows recovering T1 could contain a combination of introgressed and non-introgressed loci. Second, assuming T2 is the species tree, T1 may result from technical error while reconstructing gene trees; specifically, a paucity of variant sites that support T2 relationships (Figure 1C). This also seems likely, as the internal branch separating *C. atrocapillus* from its sister clade is very short. We recovered three rapid and successive speciation events around this node that occurred in under one million years. Finally, successive short internal branches associated with rapid speciation events could introduce high rates of ILS to the genome, possibly even rendering T2 to the anomaly zone, wherein most gene tree topologies conflict with the true species tree (Degnan and Rosenberg 2006; Cloutier et al. 2019). Each of these factors could lead to an excess of T1 across the autosomes.

### The Z-linked pseudo-autosomal region (PAR) was marked by high introgression

Our finding of decreased introgression on the Z-chromosome is consistent with Haldane’s Rule, which posits that hybrid individuals of the heterogametic sex typically exhibit lower fitness, resulting in selection against gene flow on sex chromosomes (Haldane 1922; Muller 1942; Dobzhansky 1982). Though we detected near-zero introgression on the Z-chromosome, the PAR was characterized by high introgression. This observation implies that for the Z-chromosome, Haldane’s Rule may explain baseline selection against introgression, but the elevated crossover rate at the PAR counters the impacts of linked selection in this context.

In our effort to identify the PAR, we discovered ~1.5X read depth across a swath of the Z-chromosome proximal to the PAR, likely reflecting erroneously mapped W-chromosome reads. This region of the Z-chromosome was biased toward the T4 topology, which unites the only female samples as sister species. Ideally, reads for all samples should be mapped to a female reference, which would reduce this artifact in our results because most W-reads should map correctly to the W-reference. However, our reference, *Rhea pennata,* was a male, and no female *Rhea* or tinamou chromosome-level genomes are available to our knowledge. Future studies looking at sex chromosome evolution or phylogenetics of Palaeognathes should be aware of this potential pitfall as well as any misleading effects of the PAR on phylogenetic reconstruction for sex chromosomes.

## Methods

### Sampling, assembling of whole-genomes, and variant calling

We sampled seven species of tinamous, six from an unresolved clade in the genus *Crypturellus* plus one outgroup. Specifically, we sampled *C. cinnamomeus*, *C. erythropus*, *C. atrocapillus*, *C. noctivagus*, *C. strigulosus*, and *C. undulatus*. We also sampled one outgroup taxon, *C. variegatus*. Whole genomes for these samples were previously sequenced for another study (Musher et al. 2025).

We assembled the genomes of each sample by mapping reads to a previously assembled tinamou genome in this group, *C. undulatus* (Musher, Catanach, et al. 2025). To obtain a pseudo-chromosome-level reference genome of *C. undulatus*, we used ragtag v2.1.0 (Alonge et al. 2022) to orient and align its scaffolds against the chromosome-level *Rhea pennata* genome (NCBI refseq assembly: GCF_028389875.1) using default settings. All raw data were cleaned using fastp (Chen et al. 2018) to remove adapters and low quality reads and then mapped to the *C. undulatus* reference genome using the Burrows-Wheeler algorithm (BWA) (Li and Durbin 2009). We then used bcftools (Li et al. 2009) to call Single Nucleotide Polymorphisms (SNPs) for all variants across the genome of each species, generating a Variant Call Format (VCF) file for each species. Bcftools was then used to merge the files into a single VCF, which was then filtered using vcftools v1.16 (Danecek et al. 2011) to obtain SNPs with a minimum allele frequency = 0.05, maximum missing data = 0.5, minimum quality = 30, minimum depth = 6, and maximum depth = 100.

### Genome-wide phylogenomics

We wrote custom scripts using bcftools to generate consensus sequences for 200 kb non-overlapping windows across the genome, and used vcf2phylip (https://github.com/edgardomor tiz/vcf2phylip) to convert vcf files into phylip format. We then used IQ-TREE v2.4.0 (Minh et al. 2020) to reconstruct the phylogeny for each 200kb window. In IQ-TREE, we used ModelFinder (Kalyaanamoorthy et al. 2017) to identify the best-fit substitution model among all possible models, and performed 1,000 ultrafast bootstraps across each gene tree (Hoang et al. 2018). We then used ASTRAL-III v5.7.7 (Mirarab et al. 2014; Zhang et al. 2018) to estimate the species tree from autosomal and Z-chromosome gene trees separately. Although 200 kb is a large window size, preliminary analyses suggested that smaller windows often did not produce robustly supported trees. We therefore view the “gene trees” resulting from these long windows as evidence for how phylogenetic signal varied across the genome rather than truly independent gene trees.

### Measuring genome-wide introgression

To test for putative introgression among taxa in Clade A, we first estimated *f_dM_* for 100 kb windows across the whole genome. *f_dM_* is an f-branch statistic derived from the ABBA-BABA test (D-statistic) that can quantify the magnitude of introgression assuming a species tree topology. To do so, we estimated *f_dM_* for 100 kb non-overlapping windows across the genome using the ‘ABBABABAwindows.py’ script located at https://github.com/simonhmartin/genomics_general#trees-for-sliding-windows (Martin et al. 2014). Because the recovered species tree differed between autosomal and Z-chromosome loci, we ran the *f_dM_* model twice, once assuming T1 (autosomal) as the species tree and once assuming T2 (Z-chromosome). The *f_dM_*statistic is derived from the D-statistic, which tests for introgression between one branch in a tree and two others assuming a three taxon statement with one outgroup, wherein population 1 and 2 are sister species, and population 3 is sister to the clade containing 1 and 2. Specifically, positive values of f_dM_ signify introgression between populations 2 and 3, whereas negative values signify introgression between populations 1 and 3. If there is no introgression, *f_dM_* values are expected to be symmetrically distributed around zero (Martin et al. 2014). To test for introgression assuming the topology T1, we assigned *C. atrocapillus* to population 1, *C. strigulosus* + *C. erythropus* to population 2, and *C. cinnamomeus* to population 3. To test for introgression assuming T2, we assigned *C. cinnamomeus* to population 1, *C. strigulosus* + *C. erythropus* to population 2, and *C. atrocapillus* to population 3. In both cases, we used *C. undulatus* as our outgroup population.

### Full likelihood estimation of population genetic parameters

We applied full likelihood using the MSC-M model in bpp v4.8.4 (Rannala and Yang 2003; Flouri et al. 2023) to test for gene flow between *C. atrocapillus* and *C. strigulosus* as well as estimate divergence times and effective population sizes in this clade. As most analyses indicated that T2 (the Z-chromosome tree) was the likely species tree, we used the A00 model assuming T2 as a fixed species tree. The MSC-M model assumes that alignments are derived from non-recombining segments of the genome. Therefore, for this approach, we generated new alignments and gene trees from relatively short sliding windows of 10 kb to mitigate violation of this assumption. We assigned gamma priors to tau (2,10), theta (2,100), and migration (2,1) and ran bpp on a subset of 10,000 randomly selected windows for 200,000 generations after a 10% burn-in. The MSC-M model outputs posterior distributions for mutation-scaled effective population sizes (θ) and divergence times (τ). Therefore, we assumed Ne=θ/4µ and T=τG/µ, where µ is the mutation rate, G is the generation time, and T is the divergence time in years. We assumed a mutation rate of 2.0 × 10^-8^ recovered in our previous work (Musher et al. 2025) and a generation time of 1 year (Brooks 2015).

### Identifying the pseudo-autosomal region (PAR)

PAR’s are regions of homologous recombination between heteromorphic sex chromosomes. When mapping reads to the Z-chromosome for heterogametic samples (females in ZW systems like birds), the PAR is expected to show twice the sequence coverage (read depth) compared to the remainder of the Z-chromosome (Xu et al. 2019; Wang et al. 2022). This is because the PAR is present on Z and W chromosomes (i.e., it is diploid), whereas the remaining Z-chromosome sequence is haploid in the heterogametic sex. Nevertheless, Z-chromosome regions homologous to degenerated W-chromosome regions may maintain high sequence similarity when W-degeneration has occurred relatively recently, as is the case for *Crypturellus* tinamous that have been previously studied (Wang et al. 2022). Thus, to identify putative PAR in our samples, we plotted read depth for all sites across the Z-chromosome, and defined the PAR based on the location where read depth roughly doubles to the chromosome tip in females (Xu et al. 2019).

### Modeling the distribution of genome-wide introgression

Theory predicts that introgression rates should covary with recombination rates across the genome, wherein recombination is expected to increase on the ends of macrochromosomes and on microchromosomes, but decrease on the Z-chromosome. To test these predictions, we first modeled broad patterns of introgression across the genome. First, because recombination is expected to be negatively associated with chromosome length, we estimated mean *f_dM_*for each chromosome, and modeled it against log_10_-normalized chromosome length. We then evaluated differences in *f_dM_* given chromosome type, defining macrochromosomes as those > 25 Mb in length and microchromosomes as those ≤ 25 Mb in length. We employed non-parametric Kruskal-Wallis tests for differences in rates of introgression between these categories, and followed this test with a post-hoc Dunn Test to test for individual differences between pairwise comparisons. Next, we modeled *f_dM_*as a function of the midpoint position of each 100 kb window. Here we employed a quadratic model in R to capture the expected parabolic shape of the curve, wherein introgression is expected to increase toward chromosome tips. This model was used to test the prediction that introgression should increase at the ends of macrochromosomes. We performed this model first on normalized midpoint positions for the nine macrochromosomes (Chr1–8 and ChrZ) combined, and then for Chr1–29 and ChrZ individually (note that chromosome 21, and 30–39 had too few windows to properly fit the quadratic model).

Finally, we evaluated differences in *f_dM_* between windows with different evolutionary histories as inferred in IQ-TREE (see above), again using a Kruskal-Wallis and post-hoc Dunn Test to examine differences in *f_dM_* for windows recovering each of the main topologies. We then performed logistic regression models in R v4.3.1 (R Core Team 2023) to test for a relationship between *f_dM_* and the probability of recovering each major topology (T1, T2, and T3). To evaluate the fit of these logistic regression models, we also calculated McFadden’s pseudo-R^2^, and to evaluate the effect size, we calculated the odds ratio, Ψ (the change in odds of an outcome per unit change in the predictor variable) for these models in R. Because we found a large portion of the Z-chromosome to contain mis-aligned reads, and therefore incorrectly called variants (see Results), we removed windows in this region from the analyses described in this section. Moreover, in birds, crossover breakpoints are often associated with gene promoter regions. To test for this effect here, we used a linear model to examine the relationship between gene content within 500kb of each window and *f_dM_*.

### Identifying outlier regions of introgression

As a final examination of patterns of introgression across the genome, we identified 100kb windows with measured *f_dM_* that differed significantly from the rest of the genome. To do so, we first estimated Z-scores (the number of standard deviations from the mean) for all windows. Using these Z-scores, we then calculated the p-value for each window’s *f_dM_* value using a permutation test with a Bonferroni adjustment for multiple comparisons to reduce false positives. We then looked at gene content within 500kb of all windows with *P* < 0.0001. Both Z-scores and p-values were estimated using R. Because we did not have a complete annotation for our references, we used Benchmarking of Universal Single Copy Orthologs (BUSCO) v5.3.0 (Simão et al. 2015) to annotate for 8,338 conserved avian genes. This conservative approach enables the identification of windows containing a high proportion of introgressed alleles that may represent candidates for adaptive introgression, recent introgression, or cohesion among linked co-evolved genes.

## Acknowledgements

We thank Emily Griffith and Jon Merwin for comments on an earlier version of this manuscript. Musher was funded in part by NSF grant DEB-1855812 to JDW and KPJ. TAC was partially supported by NSF grant DEB-2203228.

## Data availability

Data and scripts to replicate this research can be found at:

