## Supplemental Materials for "The genomic landscape of asymmetric introgression tracks chromosomal characteristics in tinamous (Aves: Tinamidae)"

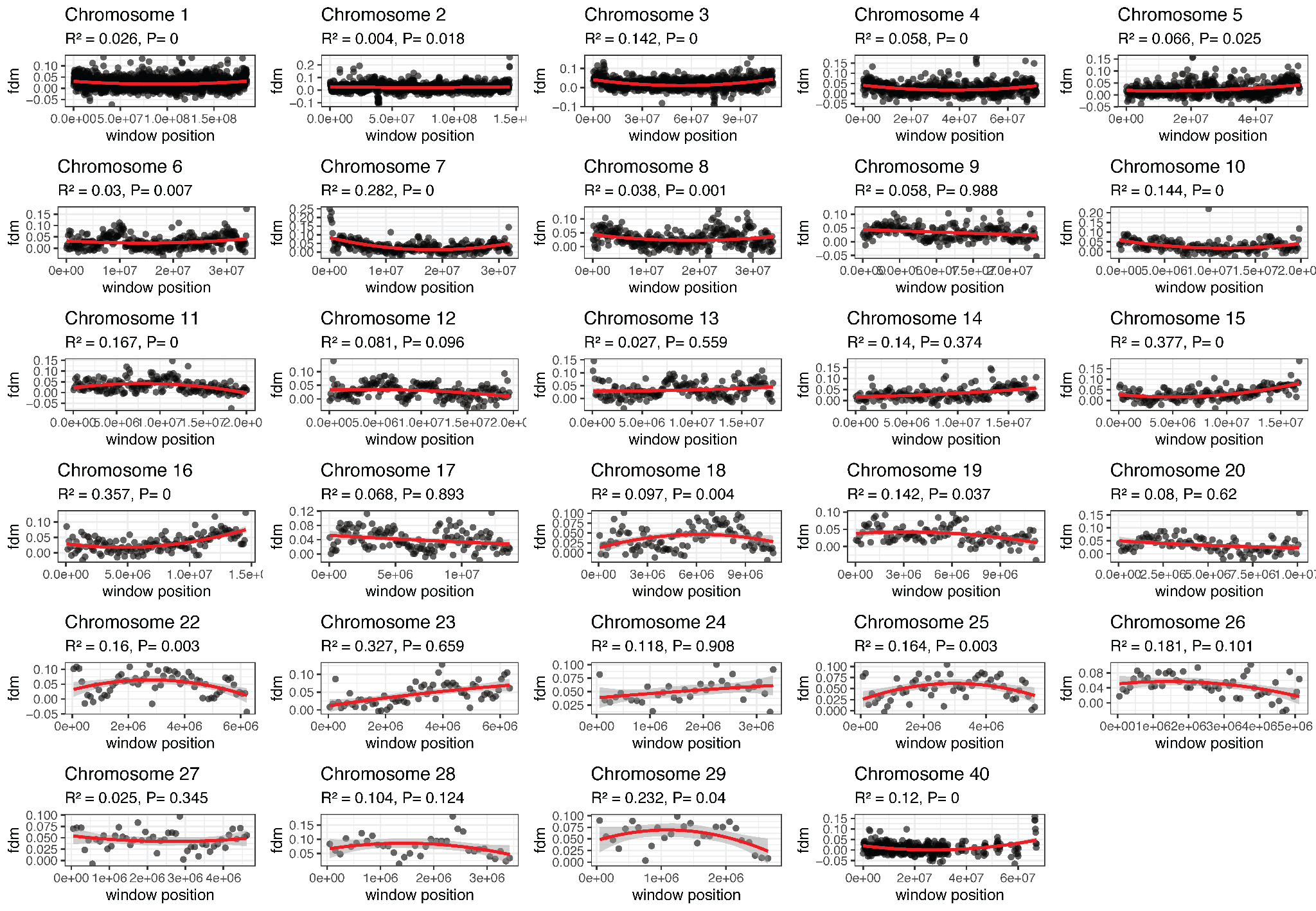

**Figure S1:** Quadratic models relating introgression position along each of the top 30 longest chromosomes in the genome using *f_dM_* results that assume T2 (the Z-chromosome topology) as the species tree. Chromosome 40 is the Z-chromosome with putatively erroneous windows removed. All macrochromosomes (Chr1–8 and Z) show significant positive quadratic relationships between position on chromosome and *f_dM_*. Note, the model for chromosome 21 failed.

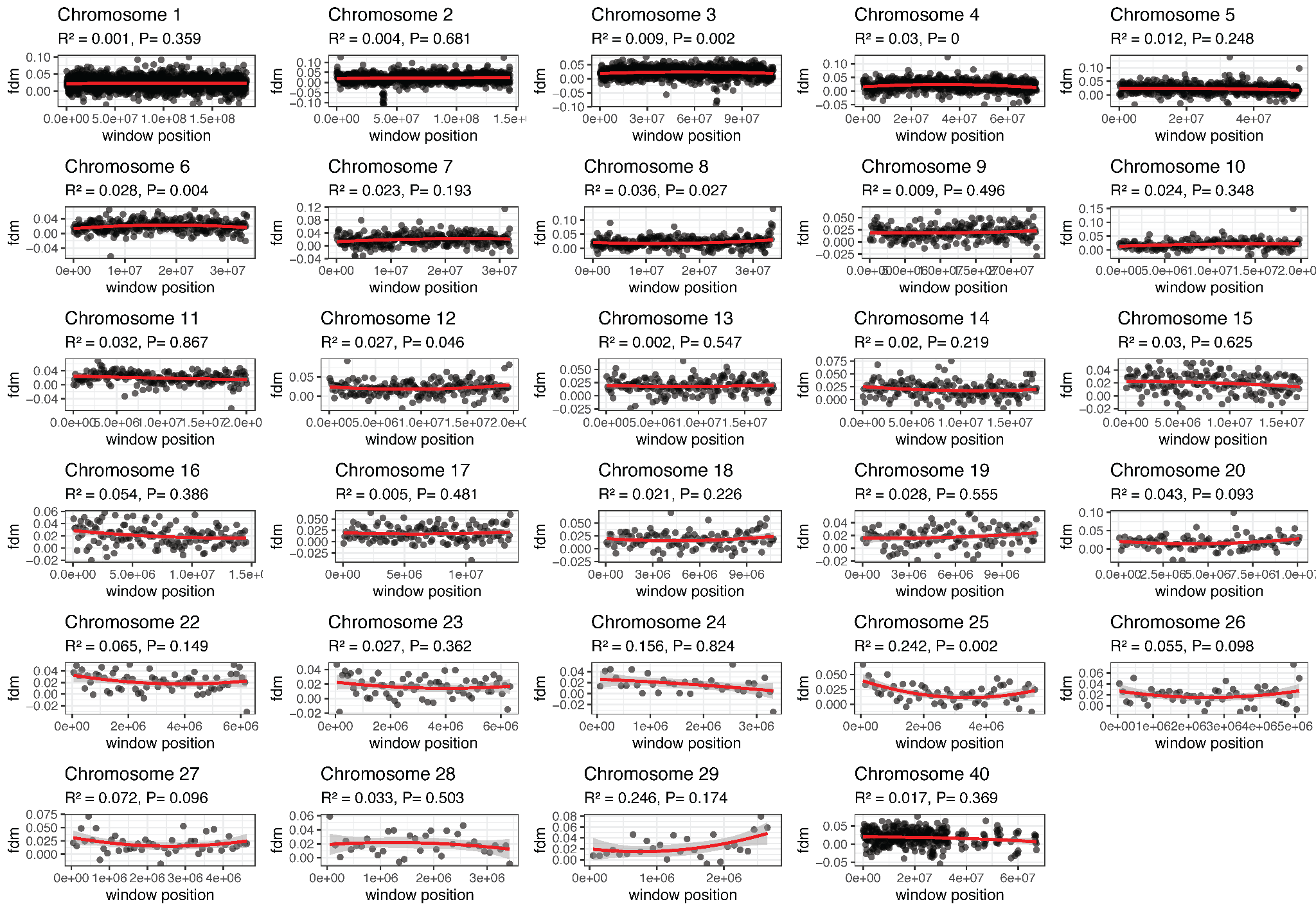

**Figure S2:** Quadratic models relating introgression position along each of the top 30 longest chromosomes in the genome using *f_dM_* results that assume T1 (the Z-chromosome topology) as the species tree. Chromosome 40 is the Z-chromosome with putatively erroneous windows removed. For most chromosomes, including macrochromosomes, there is no significant positive relationship between position on chromosome and *f_dM_*. Note, the model for chromosome 21 failed.

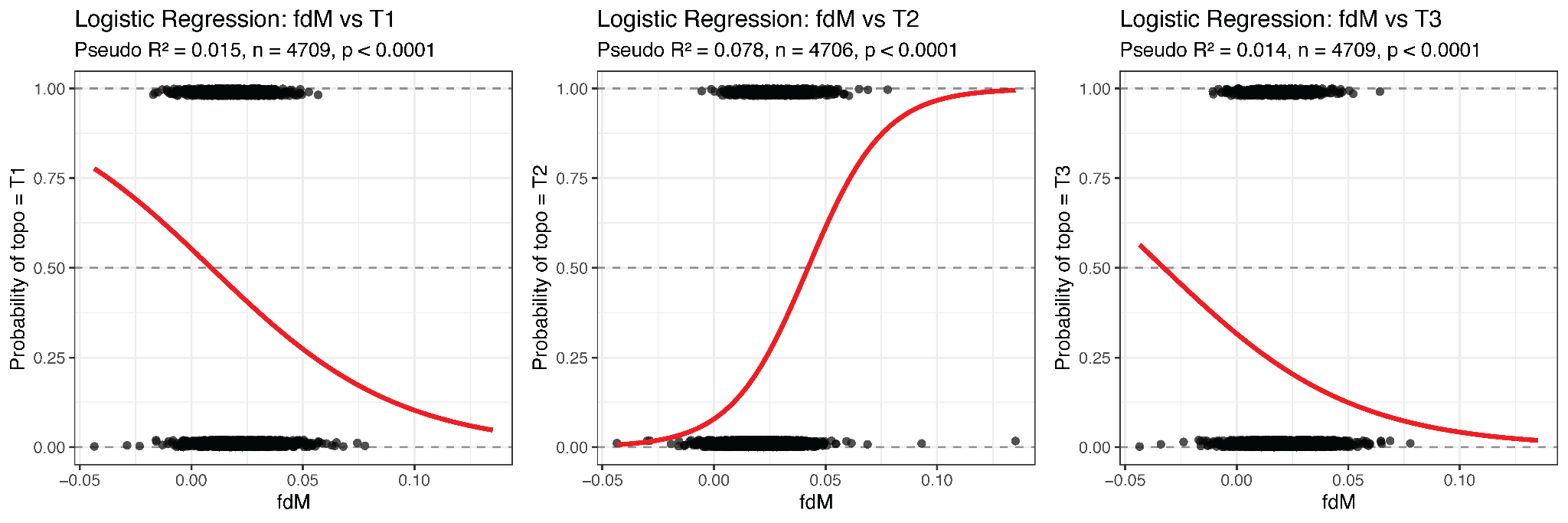

**Figure S3:** Logistic regression models (window topology ~ *f_dM_*) using *f_dM_* results that assume T1 as the species tree. Results are significant but show weak relationships

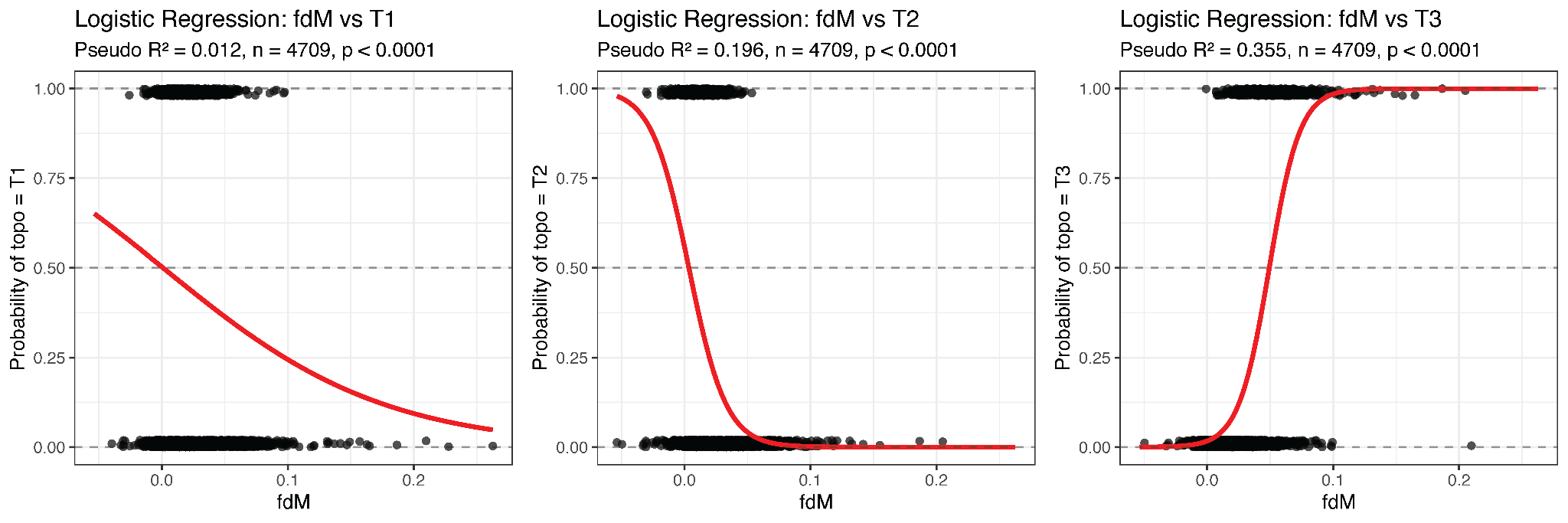

**Figure S4:** Logistic regression models (window topology ~ *f_dM_*) using *f_dM_* results that assume T2 as the species tree. Results are significant and show strong relationships for windows recovering T2 and T3.

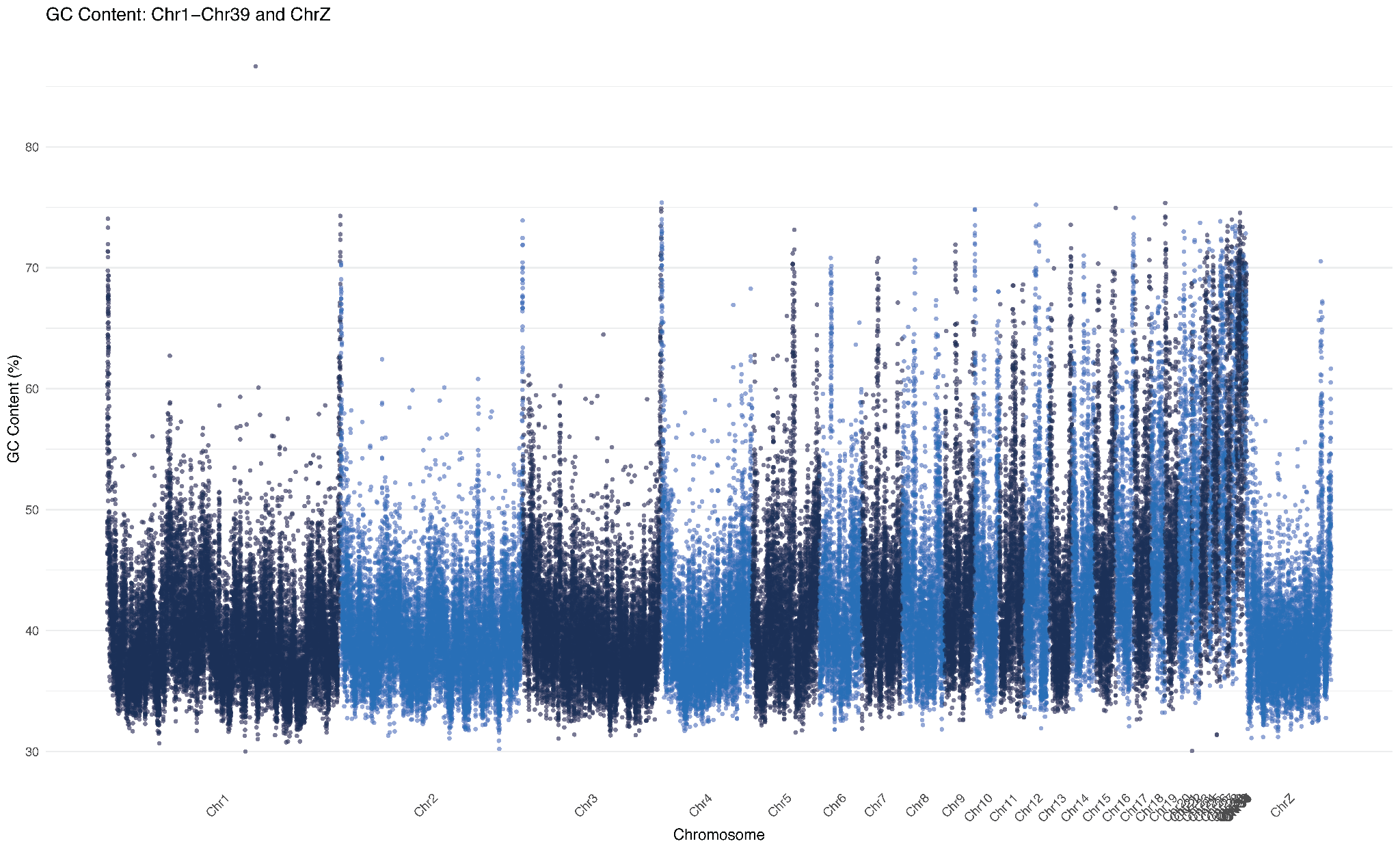

**Figure S5:** Genome-wide GC content for 10kb windows across the *C. undulatus* genome. Note the increased GC content at edges of macrochromosomes, on microchromosomes, and the PAR (end of Z-chromosome).

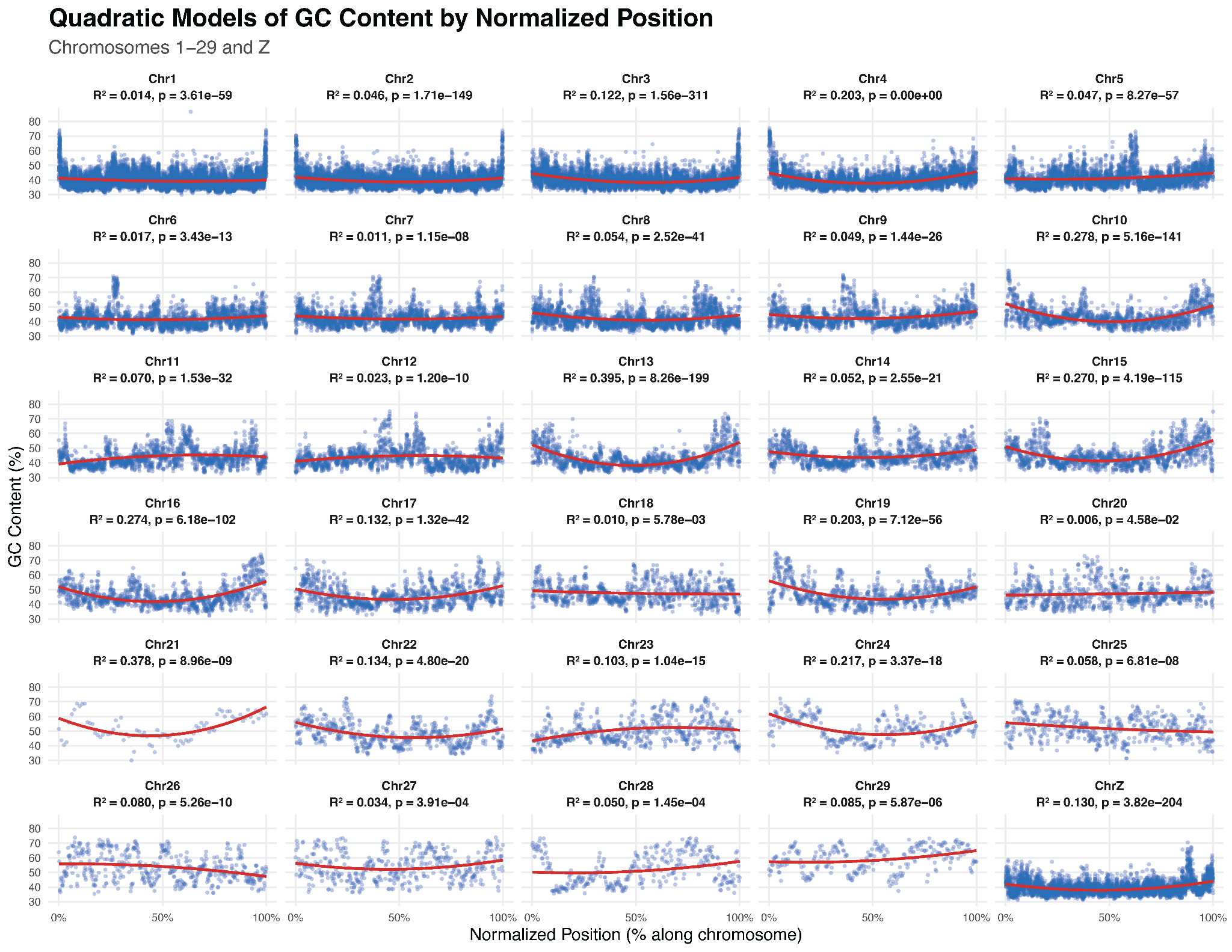

**Figure S6:** Quadratic models relating reference-genome GC content across 10kb windows to position along each of the top 30 longest chromosomes in the genome. Results mirror those in Figure S1.
